## Supplementary file for "The *Toxoplasma* monocarboxylate transporters are involved in the metabolism within the apicoplast and are linked to parasite survival"

**This Supplementary File includes:**

Figures S1-6 and Legends;

Tables S1-4.

**Figure S1-6 and Legends**

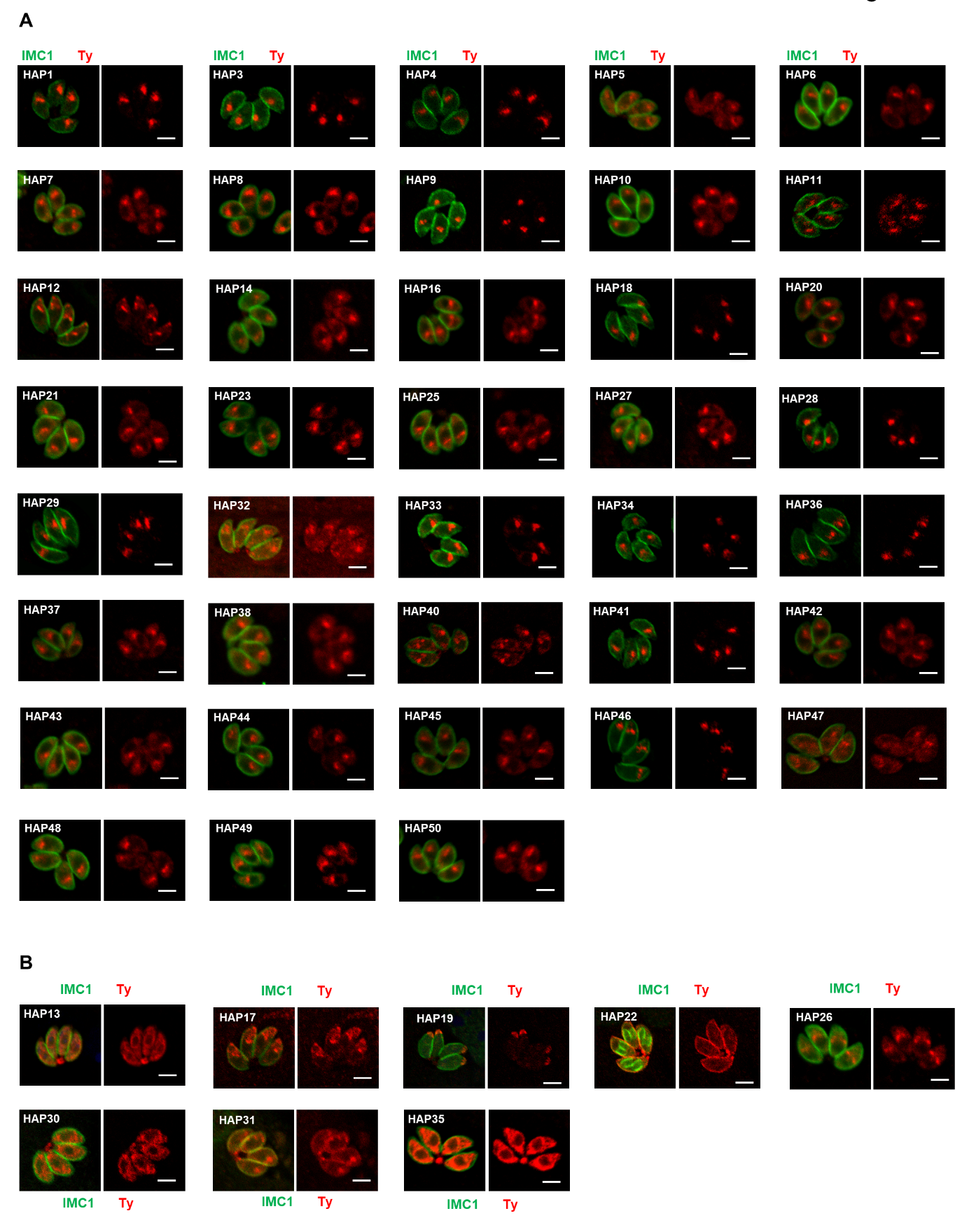

**Figure S1**. IFA analysis of HAP-6Ty fusions in parasites.

(A-B) Candidates from hyperLOPIT, ACP-BirA and APT1-TurboID were endogenously tagged with 6Ty at their C-termini using a CRISPR-Cas9 approach, followed by IFA analysis with antibodies against the inner membrane complex protein IMC1 (green) and Ty (red). Many of the HAPs were localized to a foci similar to position of the apicoplast in parasites (A), whereas others were localized to the cytosolic vesicles (e.g. HAP30, HAP13, HAP35), pellicular membrane (e.g. HAP22), or other structures (B). Scale = 5 μm. Note that HAP2, 15, 24 and 39 were not successfully tagged for IFA analysis. Three independent experiments were performed with similar outcomes and representative images are shown.

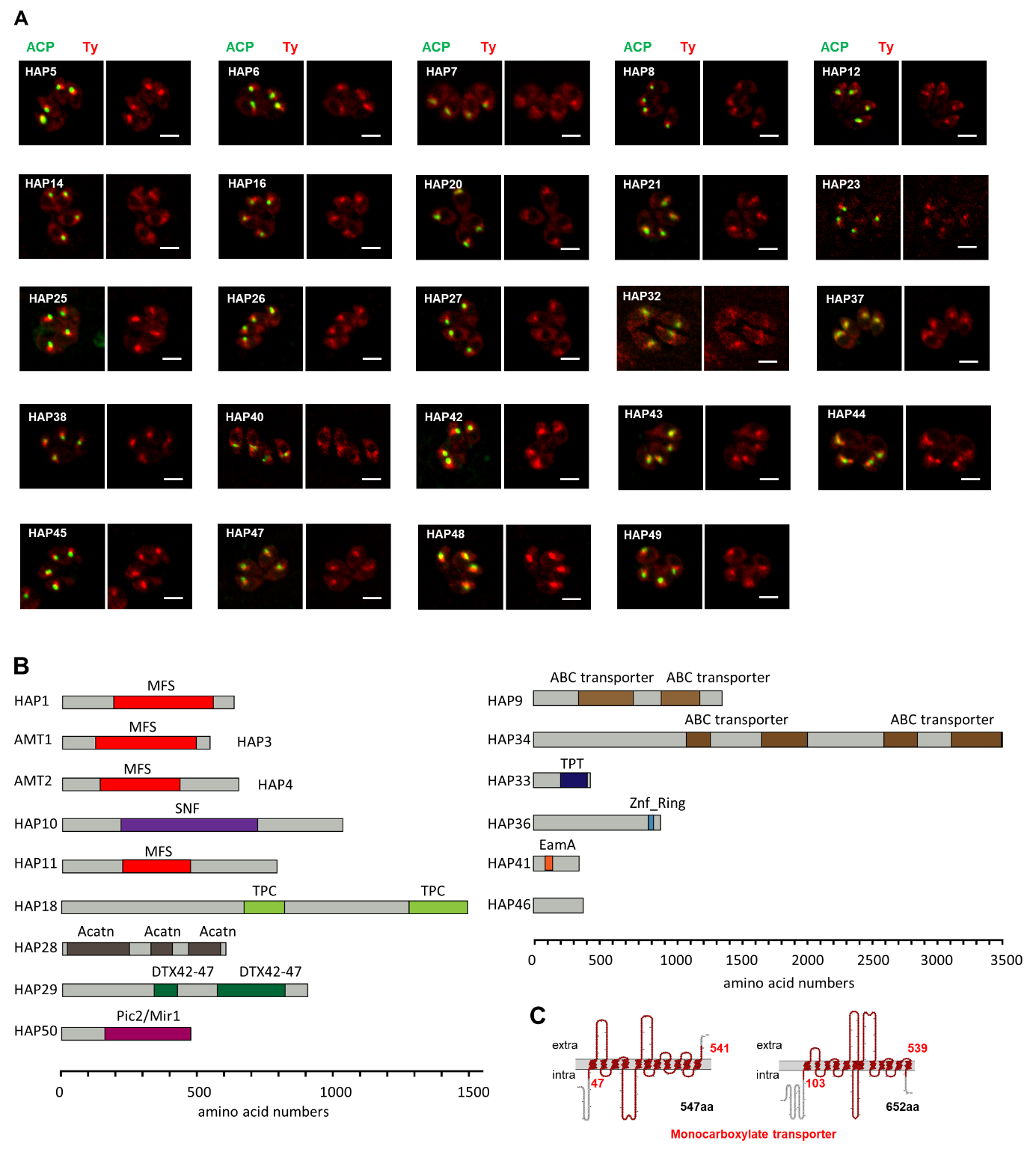

**Figure S2**. Analyses of non-apicoplast-localized HAP-fusions, and domain analysis of novel apicoplast transporters.

(A) Shown here are the HAP-Ty fusions that were not localized to the apicoplast. IFA analyses were performed with the HAP-6Ty fusions (red) and the apicoplast marker ACP (green). Scale = 5 μm.

(B) Domain analysis of newly identified apicoplast membrane proteins. Proteins were analyzed by IntroProScan (https://www.ebi.ac.uk/interpro/about/interproscan/). MFS, major facillitator transporter superfamily capable of transporting small solutes in response to chemiosmotic ion gradients (Pao et al., 1998; Walmsley et al., 1998); SNF, sodium:neutrotransmitter symporter superfamily that is responsible for exitatory amino acid transport (Malandro and Kilberg, 1996); TPC, two-pore channels that have roles in organelle integrity, inter-organelle communication and growth in *T. gondii* (Li et al., 2021); acatn, acetyl-coenzyme A transporter (Bora et al., 1999); DTX42-47, among which DTX43 is a citrate transporter responsible for loading citrate into xylem tissues and facilitate iron transport to shoots (Green and Rogers, 2004; Rogers and Guerinot, 2002); ABC, ABC transporters that are involved in export or import of a wide variety of substrates ranging from small ions to macromolecules; Pic2/Mir1-like, phosphate carriers that were reported to import copper and inorganic phosphate into mitochondria (Hamel et al., 2004; Vest et al., 2013); TPT, Sugar phosphate transporter domain with a specificity for triose phosphate (Jack et al., 2001); Znf ring, Zinc finger that may bind metals, such as iron, or no metal at all; EamA, found in many members classed as drug/metabolite transporters (Jack et al., 2001). Protein lengths were scaled by amino acid numbers (aa).

(C) The topology of two monocarboxylate transporters (AMT1 and AMT2) was analyzed by an online server PROTTER (https://wlab.ethz.ch/protter/start/).

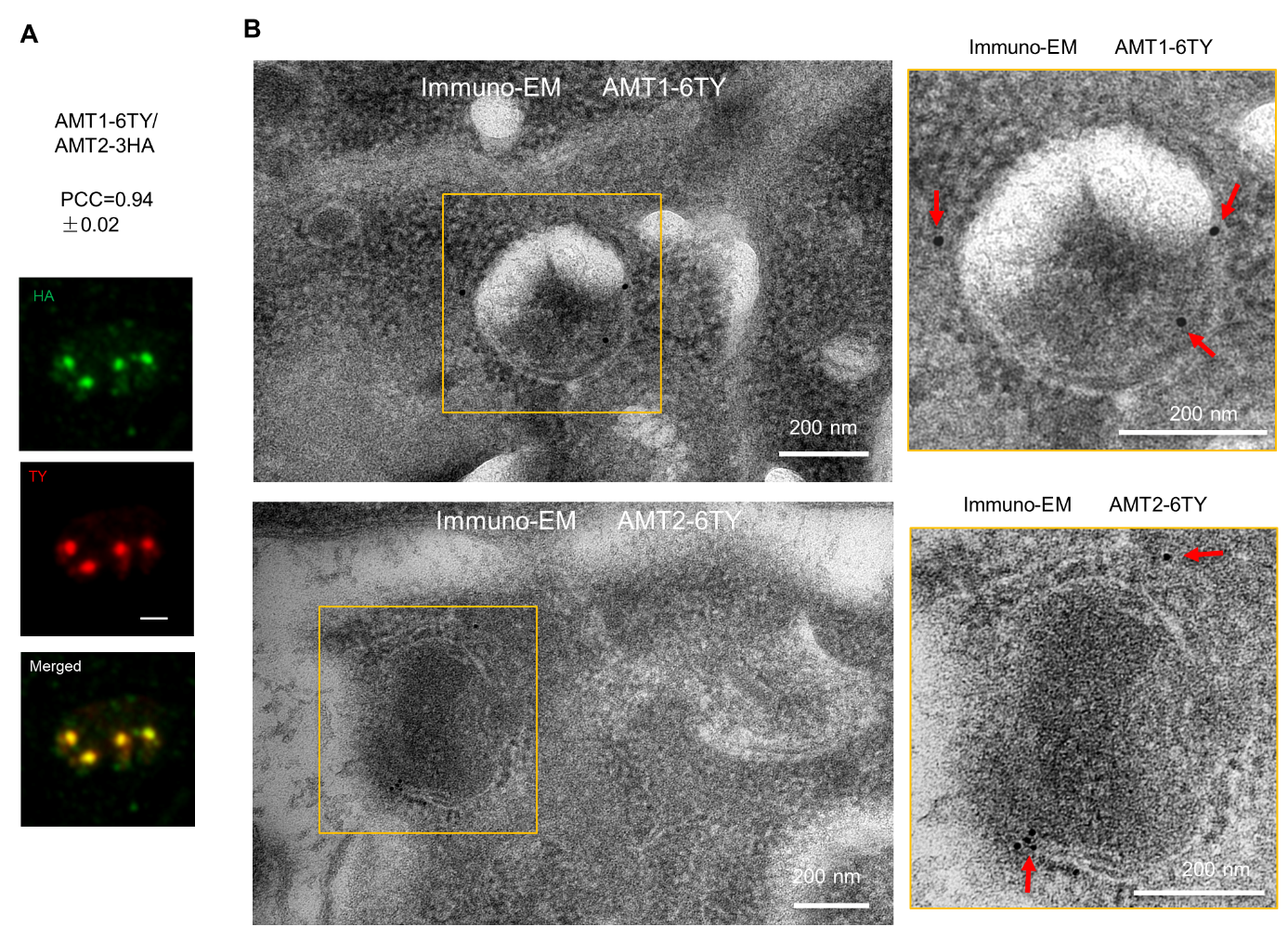

**Figure S3.** AMT1 and AMT2 are co-localized in the apicoplast

(A) Confocal images showed colocalization of AMT1-6Ty and AMT2-6HA. PCC was analyzed over the merged fluoresent foci by the NIS Elemental AR system. Data are shown with a mean ± SD (N=6).

(B) Immuno-EM of AMT1-6Ty and AMT2-6Ty. Parasites were fixed, embeded, sliced and incubated sequentially with Ty antibodies and gold particle (15 nm for AMT1 and 10 nm for AMT2) conjugated with anti-mouse antibodies. Scale bars were indicated on the images, and red arrows showed the gold particles at the apicoplast membranes.

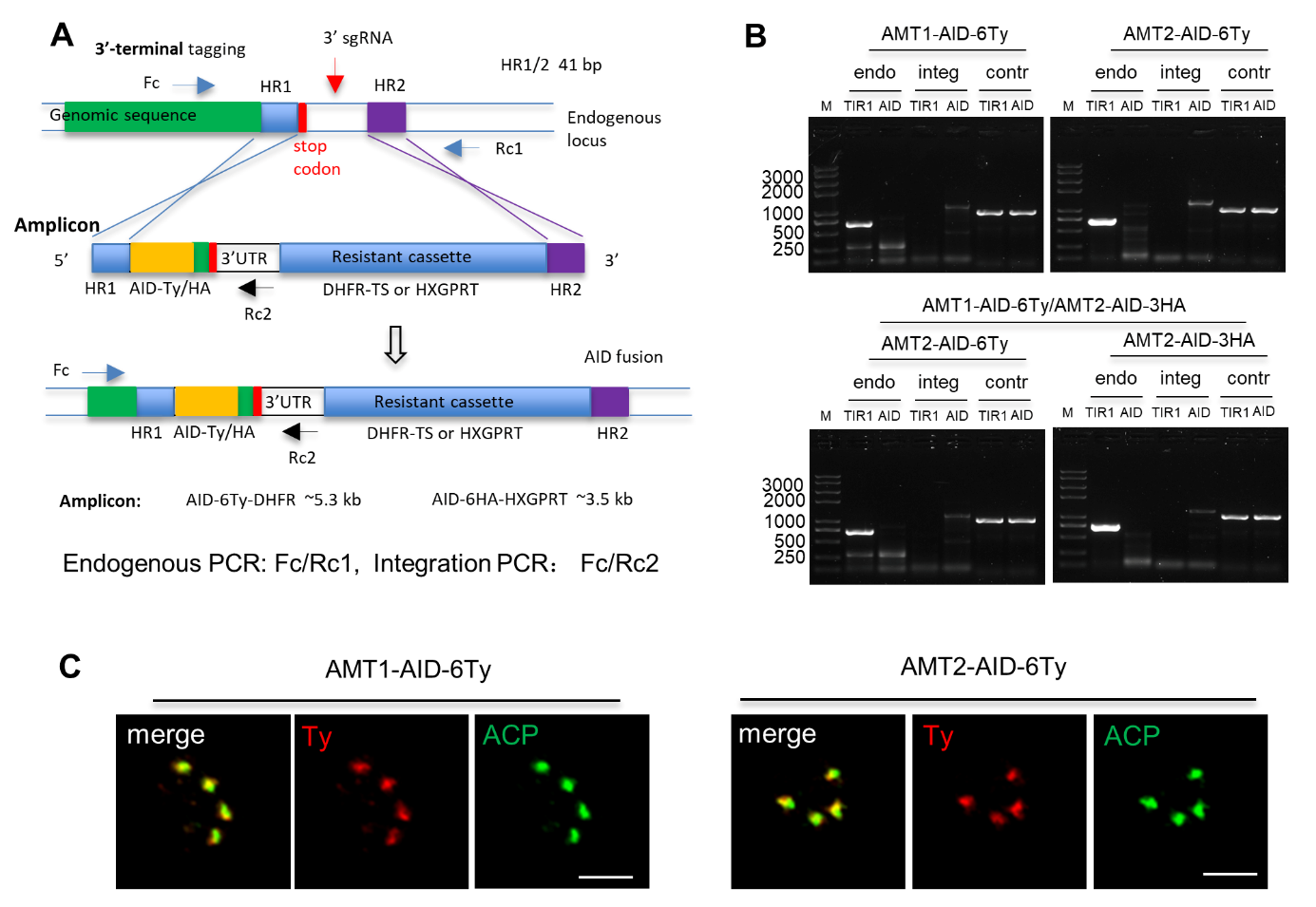

**Figure S4**. Diagnostic PCR of the AMT1-AID, AMT2-AID and dKD lines.

(A) Schematic diagram of the generation of the lines using a CRISPR-Cas9 approach. The sgRNA sequences were selected as described previously (Long et al., 2018), and integrated into a pCas9 plasmid using a DNA assembly method. The pCas9-sgRNA 3’ expresses Cas9 and sgRNA, which specifically cleave at the sgRNA region, creating a double strain DNA break (DSB) at the 3’ untranslated region of a specific gene. An amplicon containing an AID fragment, a resistant cassette, and homologous regions (HR1 and HR2) at both ends was amplified from a generic plasmid. The HR1 and HR2 targeted the amplicon to integrate at the DSB, resulting in an in-frame integration of the AID fragment at the end of the encoding sequences. The primers were designed for diagnostic PCR for testing endogenous DNA and integration of the amplicon.

(B) Diagnostic PCR of the AID lines. Endogenous (endo) and integration (integ) PCRs were performed with pairs of primers Fc/Rc (endo) and Fc/Rc2 (integ) as shown in (A), and the control (contr) PCR was performed with primers for the tubulin promoter region. The product sizes for the endo PCR in both the AID lines were 700 bp, while the sizes of the integ PCR were 1357 bp for AMT1-AID-6Ty, 1407 bp for AMT2-AID-6Ty and 1326 bp for AMT2-AID-3HA. M, DNA marker.

(C) Co-localization of AID-Ty fusions with the apicoplast marker ACP. Parasites were analyzed by IFA using antibodies against Ty (red) and ACP (green), followed by secondary antibodies conjugated with Alexa Fluors. Scale bar = 5 μm.

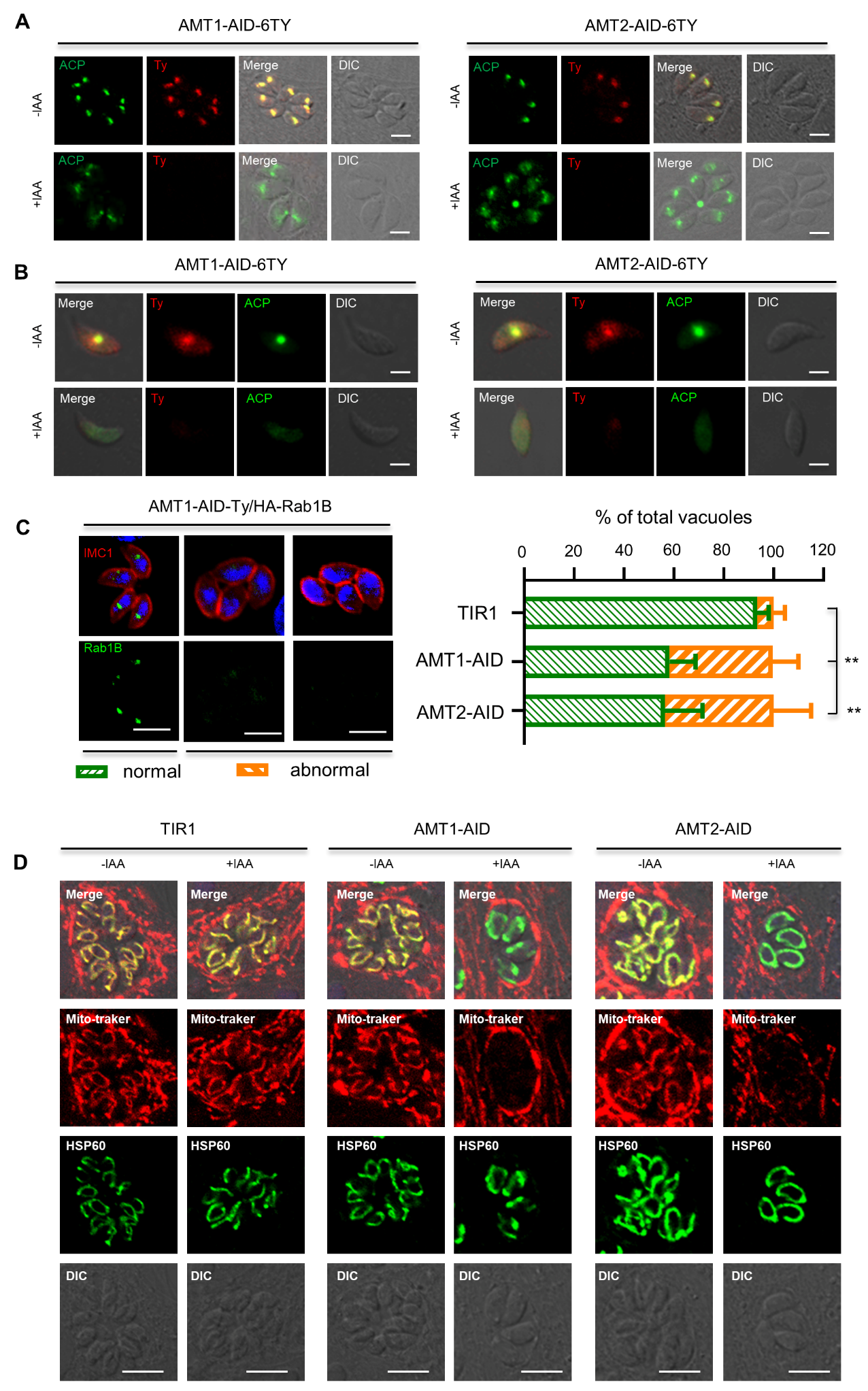

**Figure S5**. Depletion of AMT1 and AMT2 leads to apicoplast defects and loss of mitochondrial membrane potential in parasites.

(A-B) Parasites were grown in ±IAA, followed by fixation of intracellular parasites after 24 hours growth (A), or by fixation of extracellular parasites released mechanically after 24 hours growth (B). IFA was performed with antibodies against epitope tag Ty (red) and apicoplast marker ACP (green). Examples of normal ACP localization, partial diffusion of ACP in the intracellular parasites (A), and a complete diffusion of ACP in the extracellular parasites (B) were shown for the lines of AMT1-AID and AMT2-AID. Scale = 5 μm. DIC, differential interference contrast.

(C) Depletion of AMT1 and AMT2 resulted in reduced protein abundance of Rab1B in parasites. Rab1B is a key molecule that regulates endocytic trafficking of GFP vesicle, and whose abundance is regulated by protein prenylation, as demonstrated in our recent study (manuscript under review). Here, depletion of AMT1 and AMT2 resulted in similar phenotype of Rab1B, which mimics that of Rab1B in parasites with inhibited supply of isoprenoids or with knockdown of prenyl-transferase GGT-2. Parasites were grown in IAA for 12 hours and fixed for scoring of vacuoles that contain signal of normal Rab1B (wild-type) or abnormal Rab1B (reduced protein abundance), as shown in the example images. The scoring results were plotted as percentage of parasites with normal and abnormal Rab1B in the populations from three independent experiments with triplicates. Data were shown with mean ± SD and analyzed by two-way ANOVA with Tukey’s multiple comparison. **, *p*<0.005. Scale = 5 μm.

(D) Parasites grown in ±IAA for 18 hours were treated with mitotracker red for 30 min, followed by fixation of the parasites using 4% paraformaldehyde. IFA was performed with primary antibodies against mitochondrial marker HSP60 and anti-rabbit secondary antibodies conjugated with Alexa Fluor-488 (green). Scale = 5 μm.

Three independent experiments were performed with similar outcomes and representative images were shown.

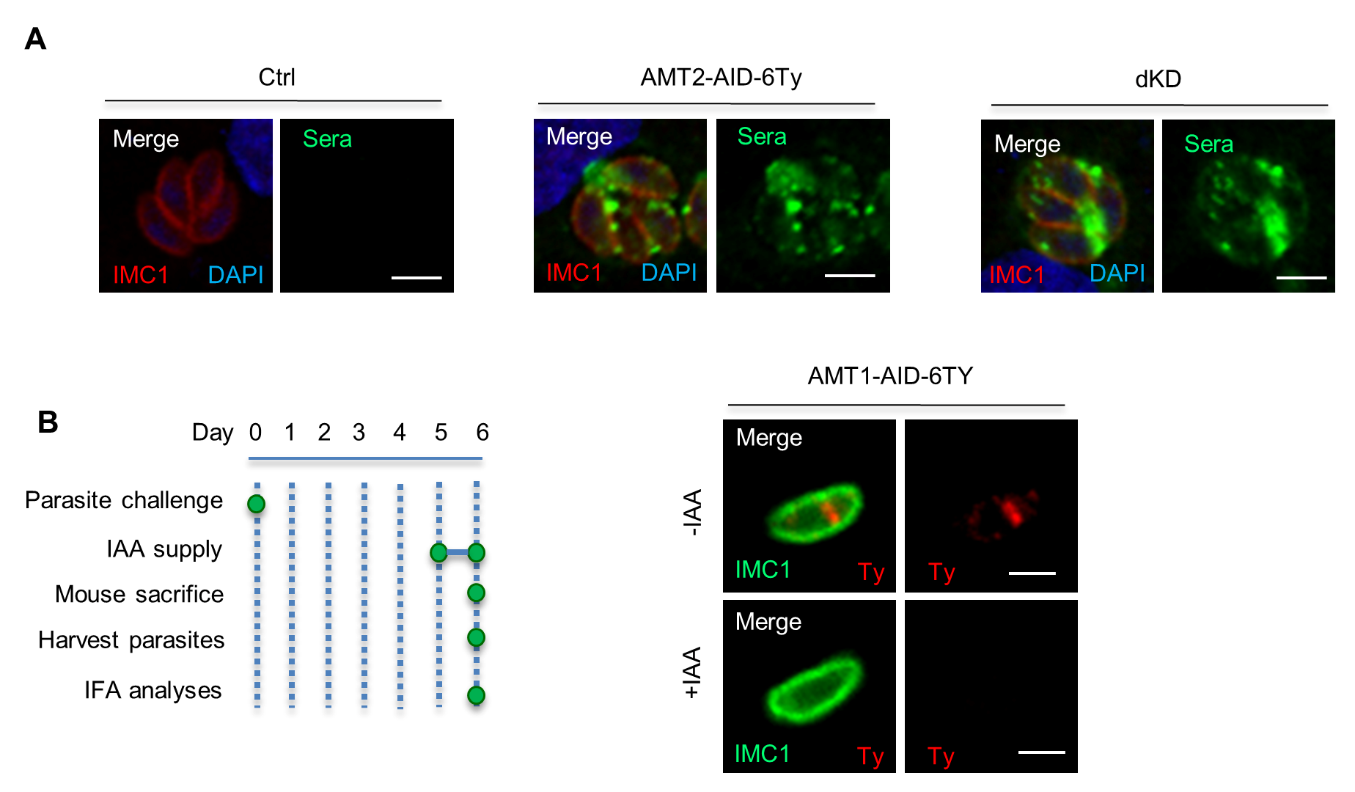

**Figure S6**. Confirmation of the mice assay by testing mice sera and protein degradation.

(A) Surviving mice were successfully infected with parasites. AMT2-AID and dKD protected mice from lethal toxoplasmosis. All surviving mice were sacrificed at day 20 for extraction of serum, followed by IFA testing of the sera at dilution ratio 1:500 using a standard protocol. Sera from blank mice (Ctrl) without *T. gondii* infection served as the IFA control. IFA results showed that the sera from surviving mice successfully recognized proteins of the parasitophorous vacuoles of analyzed *T. gondii* parasites. One representative image was shown for the mouse groups. Scale = 5 μm.

(B) The AMT1-AID fusion was efficiently degraded by IAA in mice in one day. Mice were intraperitoneally infected by parasites (100 parasites per mouse), followed by growth of parasites in mice for 5 days. The mice were randomly assigned into a control group and an induced group (n = 5 mice). The induced group was orally administered with the drinking water containing IAA for one day and intraperitoneally injected with a solution containing IAA on day 5, whereas the control group was supplied with a drinking water and an injection solution without IAA. The mice were then sacrificed on day 6 for extraction of parasites from peritoneal cavities, as demonstrated in the schematic diagram. The parasites were collected and adhered onto poly-lysine-coated coverslips for IFA, which was performed with antibodies against IMC1 (green) and Ty/HA (red). IFA showed that the AID fusion was efficiently degraded by IAA in mice within one day, and one representative image for each line infection was shown. Scale = 5 μm.

**Table S1**. Selection of candidates from previous studies of HyperLOPIT AND PfACP-BirA

| **Gene ID** | **Description** | **TMD** | **Name** | **Note** |
| --- | --- | --- | --- | --- |
| TGGT1_309580 | Transporter, MFS | 8 | HAP1 | HyperLOPIT |
| TGGT_267690 | hypothetical | 6 | HAP2 | HyperLOPIT |
| TGGT1_261070 | APT1 | 6 | APT1 | HyperLOPIT |
| TGGT1_233540 | MFS protein | 11 | HAP3 | PF3D7_0926400.1  in PfACP-BirA |
| TGGT1_297245 | MFS protein | 11 | HAP4 | PF3D7_0926400.1  in PfACP- BirA |
| TGGT1_222310 | Hypothetical  protein | 7 | HAP5 | PF3D7_0908300.2  in PfACP- BirA |
| TGGT1_249820 | ABC transporter | 4 | HAP6 | PF3D7_0302600.1  in PfACP- BirA |
| TGGT1_260310 | ABC transporter | 7 | HAP7 | PF3D7_1145500.1  in PfACP- BirA |
| TGGT1_263740 | ABC transporter | 5 | HAP8 | PF3D7_1209900.1  in PfACP- BirA |
| TGGT1_226790 | ABC  transporter | 0 | HAP9 | PF3D7_0813700.1  in PfACP- BirA |

**Table S2.** Lines used in this study.

| No. | Name | Genotype | Aim |
| --- | --- | --- | --- |
| 1 | APT1-TurboID-4Ty | RHΔ*ku80*Δ*hxgprt*; *APT1-TurboID-4Ty, DHFR-TS:DHFR* | Biotinylation |
| 2 | HAP1-6Ty | RHΔ*ku*80Δ*hxgprt; HAP1-6Ty, DHFR-TS:DHFR* | IFA |
| 3 | HAP2-6Ty | RHΔ*ku*80Δ*hxgprt; HAP2-6Ty, DHFR- TS:DHFR* | IFA |
| 4 | HAP3-6Ty（AMT1-6Ty） | RHΔ*ku*80Δ*hxgprt; HAP3-6Ty, DHFR- TS:DHFR* | IFA |
| 5 | HAP4-6Ty（AMT2-6Ty） | RHΔ*ku*80Δ*hxgprt; HAP4-6Ty, DHFR- TS:DHFR* | IFA |
| 6 | HAP5-6Ty | RHΔ*ku*80Δ*hxgprt; HAP5-6Ty, DHFR- TS:DHFR* | IFA |
| 7 | HAP6-6Ty | RHΔ*ku*80Δ*hxgprt; HAP6-6Ty, DHFR- TS:DHFR* | IFA |
| 8 | HAP7-6Ty | RHΔ*ku*80Δ*hxgprt; HAP7-6Ty, DHFR- TS:DHFR* | IFA |
| 9 | HAP8-6Ty | RHΔ*ku*80Δ*hxgprt; HAP8-6Ty, DHFR- TS:DHFR* | IFA |
| 10 | HAP9-6Ty | RHΔ*ku*80Δ*hxgprt; HAP9-6Ty, DHFR- TS:DHFR* | IFA |
| 11 | HAP10-6Ty | RHΔ*ku*80Δ*hxgprt; HAP10-6Ty, DHFR- TS:DHFR* | IFA |
| 12 | HAP11-6Ty | RHΔ*ku*80Δ*hxgprt; HAP11-6Ty, DHFR- TS:DHFR* | IFA |
| 13 | HAP12-6Ty | RHΔ*ku*80Δ*hxgprt; HAP12-6Ty, DHFR- TS:DHFR* | IFA |
| 14 | HAP13-6Ty | RHΔ*ku*80Δ*hxgprt; HAP13-6Ty, DHFR- TS:DHFR* | IFA |
| 15 | HAP14-6Ty | RHΔ*ku*80Δ*hxgprt; HAP14-6Ty, DHFR- TS:DHFR* | IFA |
| 16 | HAP15-6Ty | RHΔ*ku*80Δ*hxgprt; HAP15-6Ty, DHFR- TS:DHFR* | IFA |
| 17 | HAP16-6Ty | RHΔ*ku*80Δ*hxgprt; HAP16-6Ty, DHFR- TS:DHFR* | IFA |
| 18 | HAP17-6Ty | RHΔ*ku*80Δ*hxgprt; HAP17-6Ty, DHFR- TS:DHFR* | IFA |
| 19 | HAP18-6Ty | RHΔ*ku*80Δ*hxgprt; HAP18-6Ty, DHFR- TS:DHFR* | IFA |
| 20 | HAP19-6Ty | RHΔ*ku*80Δ*hxgprt; HAP19-6Ty, DHFR- TS:DHFR* | IFA |
| 21 | HAP20-6Ty | RHΔ*ku*80Δ*hxgprt; HAP20-6Ty, DHFR- TS:DHFR* | IFA |
| 22 | HAP21-6Ty | RHΔ*ku*80Δ*hxgprt; HAP21-6Ty, DHFR- TS:DHFR* | IFA |
| 23 | HAP22-6Ty | RHΔ*ku*80Δ*hxgprt; HAP22-6Ty, DHFR- TS:DHFR* | IFA |
| 24 | HAP23-6Ty | RHΔ*ku*80Δ*hxgprt; HAP23-6Ty, DHFR- TS:DHFR* | IFA |
| 25 | HAP24-6Ty | RHΔ*ku*80Δ*hxgprt; HAP24-6Ty, DHFR- TS:DHFR* | IFA |
| 26 | HAP25-6Ty | RHΔ*ku*80Δ*hxgprt; HAP25-6Ty, DHFR- TS:DHFR* | IFA |
| 27 | HAP26-6Ty | RHΔ*ku*80Δ*hxgprt; HAP26-6Ty, DHFR- TS:DHFR* | IFA |
| 28 | HAP27-6Ty | RHΔ*ku*80Δ*hxgprt; HAP27-6Ty, DHFR- TS:DHFR* | IFA |
| 29 | HAP28-6Ty | RHΔ*ku*80Δ*hxgprt; HAP28-6Ty, DHFR- TS:DHFR* | IFA |
| 30 | HAP29-6Ty | RHΔ*ku*80Δ*hxgprt; HAP29-6Ty, DHFR- TS:DHFR* | IFA |
| 31 | HAP30-6Ty | RHΔ*ku*80Δ*hxgprt; HAP30-6Ty, DHFR- TS:DHFR* | IFA |
| 32 | HAP31-6Ty | RHΔ*ku*80Δ*hxgprt; HAP31-6Ty, DHFR- TS:DHFR* | IFA |
| 33 | HAP32-6Ty | RHΔ*ku*80Δ*hxgprt; HAP32-6Ty, DHFR- TS:DHFR* | IFA |
| 34 | HAP33-6Ty | RHΔ*ku*80Δ*hxgprt; HAP33-6Ty, DHFR- TS:DHFR* | IFA |
| 35 | HAP34-6Ty | RHΔ*ku*80Δ*hxgprt; HAP34-6Ty, DHFR- TS:DHFR* | IFA |
| 36 | HAP35-6Ty | RHΔ*ku*80Δ*hxgprt; HAP35-6Ty, DHFR- TS:DHFR* | IFA |
| 37 | HAP36-6Ty | RHΔ*ku*80Δ*hxgprt; HAP36-6Ty, DHFR- TS:DHFR* | IFA |
| 38 | HAP37-6Ty | RHΔ*ku*80Δ*hxgprt; HAP37-6Ty, DHFR- TS:DHFR* | IFA |
| 39 | HAP38-6Ty | RHΔ*ku*80Δ*hxgprt; HAP38-6Ty, DHFR- TS:DHFR* | IFA |
| 40 | HAP39-6Ty | RHΔ*ku*80Δ*hxgprt; HAP39-6Ty, DHFR- TS:DHFR* | IFA |
| 41 | HAP40-6Ty | RHΔ*ku*80Δ*hxgprt; HAP40-6Ty, DHFR- TS:DHFR* | IFA |
| 42 | HAP41-6Ty | RHΔ*ku*80Δ*hxgprt; HAP41-6Ty, DHFR- TS:DHFR* | IFA |
| 43 | HAP42-6Ty | RHΔ*ku*80Δ*hxgprt; HAP42-6Ty, DHFR- TS:DHFR* | IFA |
| 44 | HAP43-6Ty | RHΔ*ku*80Δ*hxgprt; HAP43-6Ty, DHFR- TS:DHFR* | IFA |
| 45 | HAP44-6Ty | RHΔ*ku*80Δ*hxgprt; HAP44-6Ty, DHFR- TS:DHFR* | IFA |
| 46 | HAP45-6Ty | RHΔ*ku*80Δ*hxgprt; HAP45-6Ty, DHFR- TS:DHFR* | IFA |
| 47 | HAP46-6Ty | RHΔ*ku*80Δ*hxgprt; HAP46-6Ty, DHFR-TS:DHFR* | IFA |
| 48 | HAP47-6Ty | RHΔ*ku*80Δ*hxgprt; HAP47-6Ty, DHFR-TS:DHFR* | IFA |
| 49 | HAP48-6Ty | RHΔ*ku*80Δ*hxgprt; HAP48-6Ty, DHFR-TS:DHFR* | IFA |
| 50 | HAP49-6Ty | RHΔ*ku*80Δ*hxgprt; HAP49-6Ty, DHFR-TS:DHFR* | IFA |
| 51 | HAP50-6Ty | RHΔ*ku*80Δ*hxgprt; HAP50-6Ty, DHFR-TS:DHFR* | IFA |
| 52 | AMT1-AID | RHΔ*ku80*Δ*hxgprt; TUB1:TIR1-3FLAG, SAG1:CAT;* AMT1*-AID-6Ty, DHFR-TS:DHFR* | Conditional knockdown |
| 53 | AMT2-AID | RHΔ*ku80*Δ*hxgprt; TUB1:TIR1-3FLAG, SAG1:CAT;* AMT2*-AID-6Ty, DHFR-TS:DHFR* | Conditional knockdown |
| 54 | AMT1-AID/  AMT2-AID (dKD) | RHΔ*ku80*Δ*hxgprt; TUB1:TIR1-3FLAG, SAG1:CAT;* AMT1*-AID-6Ty, DHFR-TS:DHFR;* AMT2*-AID-3HA, DHFR-TS: HXGPRT* | Conditional knockdown |
| 55 | AMT1-AID/  PYK2-AID | RHΔ*ku80*Δ*hxgprt; TUB1:TIR1-3FLAG, SAG1:CAT;* AMT1*-AID-6Ty, DHFR-TS:DHFR; PYK*2*-AID-3HA, DHFR-TS: HXGPRT* | Conditional knockdown |
| 56 | AMT2-AID/  PYK2-AID | RHΔ*ku80*Δ*hxgprt; TUB1:TIR1-3FLAG, SAG1:CAT;* AMT2*-AID-6Ty, DHFR-TS:DHFR; PYK*2*-AID-3HA, DHFR-TS: HXGPRT* | Conditional knockdown |
| 57 | AMT1-AID/  AMT2-AID/  PYK2-AID | RHΔ*ku80*Δ*hxgprt; TUB1:TIR1-3FLAG, SAG1:CAT;* AMT1*-AID-6Ty; AMT*2*-AID-3HA; PYK*2*-AID-GFP, DHFR-TS: HXGPRT* | Conditional knockdown |
| 58 | PYK2-AID | RHΔ*ku80*Δ*hxgprt; TUB1:TIR1-3FLAG, SAG1:CAT; PYK*2*-AID-3HA, DHFR-TS: HXGPRT* | Conditional knockdown |
| 59 | AMT1-AID/APT1-6HA | RHΔ*ku80*Δ*hxgprt*; *TUB1:TIR1-3FLAG, SAG1:CAT; AMT1-AID-6Ty, DHFR-TS:DHFR; APT1-6HA, DHFR-TS-HXGPRT* | Western blot |
| 60 | AMT2-AID/APT1-6HA | RHΔ*ku80*Δ*hxgprt*; *TUB1:TIR1-3FLAG, SAG1:CAT; AMT2-AID-6Ty, DHFR-TS:DHFR; APT1-6HA, DHFR-TS-HXGPRT* | Western blot |
| 61 | AMT1-AID/AMT2-6HA | RHΔ*ku80*Δ*hxgprt*; *TUB1:TIR1-3FLAG, SAG1:CAT; AMT1-AID-6Ty, DHFR-TS:DHFR; AMT2-6HA, DHFR-TS-HXGPRT* | Western blot |
| 61 | AMT2-AID/AMT1-6HA | RHΔ*ku80*Δ*hxgprt*; *TUB1:TIR1-3FLAG, SAG1:CAT; AMT2-AID-6Ty, DHFR-TS:DHFR; AMT1-6HA, DHFR-TS-HXGPRT* | Western blot |
| 62 | AMT1-AID/ACC1-6HA | RHΔ*ku80*Δ*hxgprt*; *TUB1:TIR1-3FLAG, SAG1:CAT; AMT1-AID-6Ty, DHFR-TS:DHFR; ACC1-6HA, DHFR-TS-HXGPRT* | Western blot |
| 63 | AMT2-AID/ACC1-6HA | RHΔ*ku80*Δ*hxgprt*; *TUB1:TIR1-3FLAG, SAG1:CAT; AMT2-AID-6Ty, DHFR-TS:DHFR; ACC1-6HA, DHFR-TS-HXGPRT* | Western blot |
| 64 | AMT1-6Ty/AMT2-6HA | RHΔ*ku*80Δ*hxgprt; AMT1-6Ty, DHFR-TS:DHFR; AMT2-6HA, DHFR-TS-HXGPRT* | IFA |
| 65 | AMT1-AID/ACC1-AID | RHΔ*ku*80Δ*hxgprt; AMT1-6Ty, DHFR-TS:DHFR; ACC1-AID-3HA, DHFR-TS-HXGPRT* | Conditional knockdown |
| 66 | ACC1-AID | RHΔ*ku*80Δ*hxgprt; AMT1-6Ty, DHFR-TS:DHFR; ACC1-AID-3HA, DHFR-TS-HXGPRT* | Conditional knockdown |

**Table S3.** Plasmids used in this study.

| No. | Plasmid Name | Genotype | Application |
| --- | --- | --- | --- |
| 1 | pCas9-APT1 sgRNA 3’ | *SAG1:Cas9, U6: APT1 sgRNA 3’* | C-terminal Tagging |
| 2 | pCas9-HAP1 sgRNA 3’ | *SAG1:Cas9, U6:* HAP1 *sgRNA 3’* | C-terminal Tagging |
| 3 | pCas9-HAP2 sgRNA 3’ | *SAG1:Cas9, U6:* HAP2 *sgRNA 3’* | C-terminal Tagging |
| 4 | pCas9-HAP3 sgRNA 3’ | *SAG1:Cas9, U6:* HAP3 *sgRNA 3’* | C-terminal Tagging |
| 5 | pCas9-HAP4 sgRNA 3’ | *SAG1:Cas9, U6:* HAP4 *sgRNA 3’* | C-terminal Tagging |
| 6 | pCas9-HAP5 sgRNA 3’ | *SAG1:Cas9, U6:* HAP5 *sgRNA 3’* | C-terminal Tagging |
| 7 | pCas9-HAP6 sgRNA 3’ | *SAG1:Cas9, U6:* HAP6 *sgRNA 3’* | C-terminal Tagging |
| 8 | pCas9-HAP7 sgRNA 3’ | *SAG1:Cas9, U6:* HAP7 *sgRNA 3’* | C-terminal Tagging |
| 9 | pCas9-HAP8 sgRNA 3’ | *SAG1:Cas9, U6:* HAP8 *sgRNA 3’* | C-terminal Tagging |
| 10 | pCas9-HAP9 sgRNA 3’ | *SAG1:Cas9, U6:* HAP9 *sgRNA 3’* | C-terminal Tagging |
| 11 | pCas9-HAP10 sgRNA 3’ | *SAG1:Cas9, U6:* HAP10 *sgRNA 3’* | C-terminal Tagging |
| 12 | pCas9-HAP11 sgRNA 3’ | *SAG1:Cas9, U6:* HAP11 *sgRNA 3’* | C-terminal Tagging |
| 13 | pCas9-HAP12 sgRNA 3’ | *SAG1:Cas9, U6:* HAP12 *sgRNA 3’* | C-terminal Tagging |
| 14 | pCas9-HAP13 sgRNA 3’ | *SAG1:Cas9, U6:* HAP13 *sgRNA 3’* | C-terminal Tagging |
| 15 | pCas9-HAP14 sgRNA 3’ | *SAG1:Cas9, U6:* HAP14 *sgRNA 3’* | C-terminal Tagging |
| 16 | pCas9-HAP15 sgRNA 3’ | *SAG1:Cas9, U6:* HAP15 *sgRNA 3’* | C-terminal Tagging |
| 17 | pCas9-HAP16 sgRNA 3’ | *SAG1:Cas9, U6:* HAP16 *sgRNA 3’* | C-terminal Tagging |
| 18 | pCas9-HAP17 sgRNA 3’ | *SAG1:Cas9, U6:* HAP17 *sgRNA 3’* | C-terminal Tagging |
| 19 | pCas9-HAP18 sgRNA 3’ | *SAG1:Cas9, U6:* HAP18 *sgRNA 3’* | C-terminal Tagging |
| 20 | pCas9-HAP19 sgRNA 3’ | *SAG1:Cas9, U6:* HAP19 *sgRNA 3’* | C-terminal Tagging |
| 21 | pCas9-HAP20 sgRNA 3’ | *SAG1:Cas9, U6:* HAP20 *sgRNA 3’* | C-terminal Tagging |
| 22 | pCas9-HAP21 sgRNA 3’ | *SAG1:Cas9, U6:* HAP21 *sgRNA 3’* | C-terminal Tagging |
| 23 | pCas9-HAP22 sgRNA 3’ | *SAG1:Cas9, U6:* HAP22 *sgRNA 3’* | C-terminal Tagging |
| 24 | pCas9-HAP23 sgRNA 3’ | *SAG1:Cas9, U6:* HAP23 *sgRNA 3’* | C-terminal Tagging |
| 25 | pCas9-HAP24 sgRNA 3’ | *SAG1:Cas9, U6:* HAP24 *sgRNA 3’* | C-terminal Tagging |
| 26 | pCas9-HAP25 sgRNA 3’ | *SAG1:Cas9, U6:* HAP25 *sgRNA 3’* | C-terminal Tagging |
| 27 | pCas9-HAP26 sgRNA 3’ | *SAG1:Cas9, U6:* HAP26 *sgRNA 3’* | C-terminal Tagging |
| 28 | pCas9-HAP27 sgRNA 3’ | *SAG1:Cas9, U6:* HAP27 *sgRNA 3’* | C-terminal Tagging |
| 29 | pCas9-HAP28 sgRNA 3’ | *SAG1:Cas9, U6:* HAP28 *sgRNA 3’* | C-terminal Tagging |
| 30 | pCas9-HAP29 sgRNA 3’ | *SAG1:Cas9, U6:* HAP29 *sgRNA 3’* | C-terminal Tagging |
| 31 | pCas9-HAP30 sgRNA 3’ | *SAG1:Cas9, U6:* HAP30 *sgRNA 3’* | C-terminal Tagging |
| 32 | pCas9-HAP31 sgRNA 3’ | *SAG1:Cas9, U6:* HAP31 *sgRNA 3’* | C-terminal Tagging |
| 33 | pCas9-HAP32 sgRNA 3’ | *SAG1:Cas9, U6:* HAP32 *sgRNA 3’* | C-terminal Tagging |
| 34 | pCas9-HAP33 sgRNA 3’ | *SAG1:Cas9, U6:* HAP33 *sgRNA 3’* | C-terminal Tagging |
| 35 | pCas9-HAP34 sgRNA 3’ | *SAG1:Cas9, U6:* HAP34 *sgRNA 3’* | C-terminal Tagging |
| 36 | pCas9-HAP35 sgRNA 3’ | *SAG1:Cas9, U6:* HAP35 *sgRNA 3’* | C-terminal Tagging |
| 37 | pCas9-HAP36 sgRNA 3’ | *SAG1:Cas9, U6:* HAP36 *sgRNA 3’* | C-terminal Tagging |
| 38 | pCas9-HAP37 sgRNA 3’ | *SAG1:Cas9, U6:* HAP37 *sgRNA 3’* | C-terminal Tagging |
| 39 | pCas9-HAP38 sgRNA 3’ | *SAG1:Cas9, U6:* HAP38 *sgRNA 3’* | C-terminal Tagging |
| 40 | pCas9-HAP39 sgRNA 3’ | *SAG1:Cas9, U6:* HAP39 *sgRNA 3’* | C-terminal Tagging |
| 41 | pCas9-HAP40 sgRNA 3’ | *SAG1:Cas9, U6:* HAP40 *sgRNA 3’* | C-terminal Tagging |
| 42 | pCas9-HAP41 sgRNA 3’ | *SAG1:Cas9, U6:* HAP41 *sgRNA 3’* | C-terminal Tagging |
| 43 | pCas9-HAP42 sgRNA 3’ | *SAG1:Cas9, U6:* HAP42 *sgRNA 3’* | C-terminal Tagging |
| 44 | pCas9-HAP43 sgRNA 3’ | *SAG1:Cas9, U6:* HAP43 *sgRNA 3’* | C-terminal Tagging |
| 45 | pCas9-HAP44 sgRNA 3’ | *SAG1:Cas9, U6:* HAP44 *sgRNA 3’* | C-terminal Tagging |
| 46 | pCas9-HAP45 sgRNA 3’ | *SAG1:Cas9, U6:* HAP45 *sgRNA 3’* | C-terminal Tagging |
| 47 | pCas9-HAP46 sgRNA 3’ | *SAG1:Cas9, U6:* HAP46 *sgRNA 3’* | C-terminal Tagging |
| 48 | pCas9-HAP47 sgRNA 3’ | *SAG1:Cas9, U6:* HAP47 *sgRNA 3’* | C-terminal Tagging |
| 49 | pCas9-HAP48 sgRNA 3’ | *SAG1:Cas9, U6:* HAP48 *sgRNA 3’* | C-terminal Tagging |
| 50 | pCas9-HAP49 sgRNA 3’ | *SAG1:Cas9, U6:* HAP49 *sgRNA 3’* | C-terminal Tagging |
| 51 | pCas9-HAP50 sgRNA 3’ | *SAG1:Cas9, U6:* HAP50 *sgRNA 3’* | C-terminal Tagging |
| 52 | pCas9-PYK2 sgRNA 3’ | *SAG1:Cas9, U6: PYK2 sgRNA 3’* | C-terminal Tagging |
| 53 | pCas9-ACC1 sgRNA 3’ | *SAG1:Cas9, U6: ACC1 sgRNA 3’* | C-terminal Tagging |
| 54 | pLinker-TurboID-4Ty-DHFR-LoxP | *Linker-TurboID-4Ty, LoxP-DHFR-TS:DHFR-LoxP* | C-terminal Tagging |
| 55 | pLinker-6Ty-DHFR-LoxP | *Linker-6Ty, LoxP-DHFR-TS:DHFR-LoxP* | C-terminal Tagging |
| 56 | pLinker-6HA-HXGPRT-LoxP | *Linker-6HA, LoxP-DHFR-TS:HXGPRT-LoxP* | C-terminal Tagging |
| 57 | pLinker-AID-6Ty-DHFR-LoxP | *Linker-AID-6Ty, LoxP-DHFR-TS:DHFR-LoxP* | C-terminal Tagging |
| 58 | pLinker-AID-3HA-HXGPRT-LoxP | *Linker-AID-3HA, LoxP-DHFR-TS:HXGPRT-LoxP* | C-terminal Tagging |
| 59 | pLinker-AID-GFP-DHFR-LoxP | *Linker-AID-GFP, LoxP-DHFR-TS:DHFR-LoxP* | C-terminal Tagging |
| 60 | pET28a-AMT1-HA-AMP | *T7 promoter-AMT1-HA, AMPR promoter-AMP* | Heterologous expression |
| 61 | pET28a-AMT2-Ty-Kan | *T7 promoter-AMT2-Ty, AMPR promoter-Kan* | Heterologous expression |

**Table S4.** Primers used in this study.

|  | Templates | Primer pairs and sequence(5’→3‘) | Fragments |
| --- | --- | --- | --- |
| TGGT1_261070  (APT1) | pCas9 | TAAATGGGGATGTCAAGTT gactgggctgttgagaaacg GTTTTAGAGCTAGAAATAGC | Specific sgRNA |
|  | Generic tagging plasmids | L: gcaggcaccctgatctattctctctcgaagaccaagtacgga AGGGCGAATTGGAGCTC  T: tccctcgcttctctccccctccccgactgggctgttgagaa TCCCAGTCACGACGTTG | Endogenous Tagging fragments |
| TGGT1_309580  (HAP1) | pCas9 | TAAATGGGGATGTCAAGTT gcagaactcccagataccca GTTTTAGAGCTAGAAATAGC | Specific sgRNA |
|  | Generic tagging plasmids | L: tcgagatcgaggtgtccgatggagaggcaaactgccctgcc gctagcAAGGGCTCGGG  T: tgcctctcgaggagtctcgcattcacagcgcccagccctgg TAGGGCGAATTGGAGCTC | Endogenous Tagging fragments |
| TGGT1_267690  (HAP2) | pCas9 | TAAATGGGGATGTCAAGTT cttcgatgctttgatgtccg GTTTTAGAGCTAGAAATAGC | Specific sgRNA |
|  | Generic tagging plasmids | L: tttacgacgttcactggtatcttcgctggtcttcgatgctt GCTAGCAAGGGCTCGG  T: ccatccgcgggcagggtcagctcgcttctcgctagcctcgg TAGGGCGAATTGGAGCTC | Endogenous Tagging fragments |
| TGGT1_233540  (HAP3/AMT1) | pCas9 | TAAATGGGGATGTCAAGTT ggcgggacgtttcttagatg GTTTTAGAGCTAGAAATAGC | Specific sgRNA |
|  | Generic tagging plasmids | L: ctccgacacccaggaaacgacactcttttcagaaacagtca GCTAGCAAGGGCTCGG  T: attttgtcgtgattgtttctgaaaggcgggacgtttcttag TAGGGCGAATTGGAGCTC | Endogenous Tagging fragments |
|  | *T. gondii* DNA | Fc: TTCTGCGACTCCGCGTAC  Rc1: CCTTCACGACAGACACCC  Rc2: CACTTCTCGTACTATGGCCGG | Diagnostic PCR |
| TGGT1_297245  (HAP4/AMT2) | pCas9 | TAAATGGGGATGTCAAGTT gcgcaagtcatacgctcgtg GTTTTAGAGCTAGAAATAGC | Specific sgRNA |
|  | Generic tagging plasmids | L: caacagacgcgctgccgtaccgcttcccgacgtactcgccc GCTAGCAAGGGCTCGG  T: agcgaaagtcgctccacgagagagacactgattctccacac TAGGGCGAATTGGAGCTC | Endogenous Tagging fragments |
|  | *T. gondii* DNA | Fc: GTGTGTGTGTATGCACCC  Rc1: CACTTTCGGGCATTGCAT  Rc2: CACTTCTCGTACTATGGCCGG | Diagnostic PCR |
| TGGT1_222310  (HAP5) | pCas9 | TAAATGGGGATGTCAAGTT gccgtttccagacacaacgc GTTTTAGAGCTAGAAATAGC | Specific sgRNA |
|  | Generic tagging plasmids | L: acggcgcgtattaccagcagtactatcctcctcctgctctg GCTAGCAAGGGCTCGG  T: accgaaatttgcaaaaaaccagatccacaccttggccagcg TAGGGCGAATTGGAGCTC | Endogenous Tagging fragments |
| TGGT1_249820  (HAP6) | pCas9 | TAAATGGGGATGTCAAGTT acatgactgcgagactctgt GTTTTAGAGCTAGAAATAGC | Specific sgRNA |
|  | Generic tagging plasmids | L: atgctcaactttttgccgacgaaaaaccgaaggaatctgtt GCTAGCAAGGGCTCGG  T: gtttctgttcctttggtgttcacaccgtttgaagacccaca TAGGGCGAATTGGAGCTC | Endogenous Tagging fragments |
| TGGT1_260310  (HAP7) | pCas9 | TAAATGGGGATGTCAAGTT ggagacgaaacgtccccctg GTTTTAGAGCTAGAAATAGC | Specific sgRNA |
|  | Generic tagging plasmids | L: gggtgtacagacagctcgtgaacatgtcgcgttactcggcg GCTAGCAAGGGCTCGG  T: ctttcggccgtagcgcctctttcgctgaggagacgaaacgt TAGGGCGAATTGGAGCTC | Endogenous Tagging fragments |
| TGGT1_263740  (HAP8) | pCas9 | TAAATGGGGATGTCAAGTT gagcgaacgcctctcagctt GTTTTAGAGCTAGAAATAGC | Specific sgRNA |
|  | Generic tagging plasmids | L: cttcttctctgtgcagcctcgactgcaatgcctcctgcaac GCTAGCAAGGGCTCGG  T: agtggttgagagttttttctcatgaagagcgaacgcctctc TAGGGCGAATTGGAGCTC | Endogenous Tagging fragments |
| TGGT1_226790  (HAP9) | pCas9 | TAAATGGGGATGTCAAGTT gggctgccagttacgttgtg GTTTTAGAGCTAGAAATAGC | Specific sgRNA |
|  | Generic tagging plasmids | L: gaagagtcaaaggtgtaaagaatttgaagaggtggacagag GCTAGCAAGGGCTCGG  T: tggacccggagaaaaccaagcgaatccaggcaaccccccac TAGGGCGAATTGGAGCTC | Endogenous Tagging fragments |
| TGGT1_264870  (HAP10) | pCas9 | TAAATGGGGATGTCAAGTT agtggcgagaggcgaaacaa GTTTTAGAGCTAGAAATAGC | Specific sgRNA |
|  | Generic tagging plasmids | L: ccagcgggatcgcagggaaacaagacacagagagaagcaac GCTAGCAAGGGCTCGG  T: ctcctccgctcttgcggttcctctctctctctcttcccttg TAGGGCGAATTGGAGCTC | Endogenous Tagging fragments |
| TGGT1_262660  (HAP11) | pCas9 | TAAATGGGGATGTCAAGTT acatgtacagccgttgggtc GTTTTAGAGCTAGAAATAGC | Specific sgRNA |
|  | Generic tagging plasmids | L: ctgcttttcttgtaacgcaggttagacgtcttgaatatggg GCTAGCAAGGGCTCGG  T: caccctccaaaagccgtttcgagcacatgtacagccgttgg TAGGGCGAATTGGAGCTC | Endogenous Tagging fragments |
| TGGT1_226020  (HAP12) | pCas9 | TAAATGGGGATGTCAAGTT agacgagggctggtcgcgcc GTTTTAGAGCTAGAAATAGC | Specific sgRNA |
|  | Generic tagging plasmids | L: ccagctcacatggcgaccgtagtgggatcagatacgttccg GCTAGCAAGGGCTCGG  T: tttagttctgacatagacaacagaagacgagggctggtcgc TAGGGCGAATTGGAGCTC | Endogenous Tagging fragments |
| TGGT1_320020  (HAP13) | pCas9 | TAAATGGGGATGTCAAGTT acacgaggatacagcatcac GTTTTAGAGCTAGAAATAGC | Specific sgRNA |
|  | Generic tagging plasmids | L: cggacgaagacccgcgtgcaccgagtccagaggtcgctgtg GCTAGCAAGGGCTCGGGC  T: ggcaaaaaaatcgaatcattctcggtccctagactcctgtg TAGGGCGAATTGGAGCTC | Endogenous Tagging fragments |
| TGGT1_280400  (HAP14) | pCas9 | TAAATGGGGATGTCAAGTT aagcggcgagaagatcagtt GTTTTAGAGCTAGAAATAGC | Specific sgRNA |
|  | Generic tagging plasmids | L: ttcgcgctgtctcgctcttcatcaataccctccttctttcc GCTAGCAAGGGCTCGG  T: aaaaaggaacaaacatacggtgaaggaagcggcgagaagat TAGGGCGAATTGGAGCTC | Endogenous Tagging fragments |
| TGGT1_289070  (HAP15) | pCas9 | TAAATGGGGATGTCAAGTT ggaatgctgcgtcaggccct GTTTTAGAGCTAGAAATAGC | Specific sgRNA |
|  | Generic tagging plasmids | L: ctcacacaaacagtcctggagagtctgataatctcatggttt GCTAGCAAGGGCTCGG  T: gttccactttgagctcgacctgccgcttccgctcgcctagg TAGGGCGAATTGGAGCTC | Endogenous Tagging fragments |
| TGGT1_252340  (HAP16) | pCas9 | TAAATGGGGATGTCAAGTT atacctcacttcgagatgcg GTTTTAGAGCTAGAAATAGC | Specific sgRNA |
|  | Generic tagging plasmids | L: tcctcggaacctgcaacctcgtcggctactcttacggtatt GCTAGCAAGGGCTCGG  T: aagtcgaagttgctctcctgcaaatgtctgtgtgaccacgc TAGGGCGAATTGGAGCTC | Endogenous Tagging fragments |
| TGGT1_216820  (HAP17) | pCas9 | TAAATGGGGATGTCAAGTT ggctgaacgacagccgttgt GTTTTAGAGCTAGAAATAGC | Specific sgRNA |
|  | Generic tagging plasmids | L: ctactgcggatagcgataatgaaacacatcggagagtggcg GCTAGCAAGGGCTCGG  T: cctgctcagtactgctgccagacacccagcaaagaccaaca TAGGGCGAATTGGAGCTC | Endogenous Tagging fragments |
| TGGT1_311080  (HAP18) | pCas9 | TAAATGGGGATGTCAAGTT gaggcttgacgctcctgtga GTTTTAGAGCTAGAAATAGC | Specific sgRNA |
|  | Generic tagging plasmids | L: aggtggcttcgtcaagcagcagttcttggcgtcctgtgctc GCTAGCAAGGGCTCGG  T: gaggcacagagcgatgcgtctggcgaggcttgacgctcctg TAGGGCGAATTGGAGCTC | Endogenous Tagging fragments |
| TGGT1_238390  (HAP19) | pCas9 | TAAATGGGGATGTCAAGTT acagagaatctcctctctcg GTTTTAGAGCTAGAAATAGC | Specific sgRNA |
|  | Generic tagging plasmids | L: tgtctatgggctctggccaccttgtcggcgatcgtagcgcg GCTAGCAAGGGCTCGG  T: gtcggtcgaggtctttcgctccatcagttctctgtccccga TAGGGCGAATTGGAGCTC | Endogenous Tagging fragments |
| TGGT1_312622  (HAP20) | pCas9 | TAAATGGGGATGTCAAGTT agtgtctccccttcgcgatg GTTTTAGAGCTAGAAATAGC | Specific sgRNA |
|  | Generic tagging plasmids | L: ctcagcacgcgcagacctctggaaaggcgtcgcatggcctt GCTAGCAAGGGCTCGG  T: aacgttctatttcaagtgccaagagagtgtctccccttcgc TAGGGCGAATTGGAGCTC | Endogenous Tagging fragments |
| TGGT1_283730  (HAP21) | pCas9 | TAAATGGGGATGTCAAGTT agtctgcagtcgctcgcggt GTTTTAGAGCTAGAAATAGC | Specific sgRNA |
|  | Generic tagging plasmids | L: tcctgttcgtgaagtacatttattctcgcatcaagtcggac GCTAGCAAGGGCTCGG  T: ccgcgccagaccttcttccgggccaagtctgcagtcgctcg TAGGGCGAATTGGAGCTC | Endogenous Tagging fragments |
| TGGT1_249580  (HAP22) | pCas9 | TAAATGGGGATGTCAAGTT agattgccgggatcggattt GTTTTAGAGCTAGAAATAGC | Specific sgRNA |
|  | Generic tagging plasmids | L: gagcggtagagtgtagtggagacggcacagcaccagacggc GCTAGCAAGGGCTCGG  T: aattttcttgttctctgccctggccttccgggcacccgaaa TAGGGCGAATTGGAGCTC | Endogenous Tagging fragments |
| TGGT1_270200  (HAP23) | pCas9 | TAAATGGGGATGTCAAGTT attacagaaggccagtgtac GTTTTAGAGCTAGAAATAGC | Specific sgRNA |
|  | Generic tagging plasmids | L: gagcagatgcagatcttatttggggaaagcgtagaaagcag GCTAGCAAGGGCTCGG  T: cgtgtgctgctctacacaactgaaacggcggaagtcctgta TAGGGCGAATTGGAGCTC | Endogenous Tagging fragments |
| TGGT1_263690  (HAP24) | pCas9 | TAAATGGGGATGTCAAGTT acaccactgagttgtgtagt GTTTTAGAGCTAGAAATAGC | Specific sgRNA |
|  | Generic tagging plasmids | L: cgtttatcgacagtgccggacgcgttgctcgcttcatggtt GCTAGCAAGGGCTCGG  T: tgtggaaggggcgctcgagtcgagacaccactgagttgtgt TAGGGCGAATTGGAGCTC | Endogenous Tagging fragments |
| TGGT1_261430  (HAP25) | pCas9 | TAAATGGGGATGTCAAGTT gcactcacacgccagggcgc GTTTTAGAGCTAGAAATAGC | Specific sgRNA |
|  | Generic tagging plasmids | L: tgcaggagtggaacctcgtcgacgatgatttgagcgacgag GCTAGCAAGGGCTCGG  T: gctcggtctgcgtttgtttcctcttgcactcacacgccagg TAGGGCGAATTGGAGCTC | Endogenous Tagging fragments |
| TGGT1_223600  (HAP26) | pCas9 | TAAATGGGGATGTCAAGTT gttcgagcgacgaaaagcaa GTTTTAGAGCTAGAAATAGC | Specific sgRNA |
|  | Generic tagging plasmids | L: gaggcctgcagcagctgtcgatttgggacgtccacacctcc GCTAGCAAGGGCTCGG  T: acggtcaaaaggcagtcaacaacgagttcgagcgacgaaaa TAGGGCGAATTGGAGCTC | Endogenous Tagging fragments |
| TGGT1_310140  (HAP27) | pCas9 | TAAATGGGGATGTCAAGTT gttgaagaggaagcgaacct GTTTTAGAGCTAGAAATAGC | Specific sgRNA |
|  | Generic tagging plasmids | L: aaacgctcaagctgctggcgacctgcgacgaggactctgtg GCTAGCAAGGGCTCGG  T: cgagtgagactttgaaacgtttttcccagtctcttccgagg TAGGGCGAATTGGAGCTC | Endogenous Tagging fragments |
| TGGT1_215940  (HAP28) | pCas9 | TAAATGGGGATGTCAAGTT gatgcgcgcgagacgtgact GTTTTAGAGCTAGAAATAGC | Specific sgRNA |
|  | Generic tagging plasmids | L: gaggcgactccgacggaaccgagctgaagagagtggagaag GCTAGCAAGGGCTCGG  T: tttccgcgttgctctcccctttctctcgttttcttcctagt TAGGGCGAATTGGAGCTC | Endogenous Tagging fragments |
| TGGT1_291940  (HAP29) | pCas9 | TAAATGGGGATGTCAAGTT aagcgtttccagccgccccg GTTTTAGAGCTAGAAATAGC | Specific sgRNA |
|  | Generic tagging plasmids | L: accgtctggctgtccaacacagtaaaggagatgccttgtgt GCTAGCAAGGGCTCGG  T: ggtttcgccaagatcaaacaagtttcccactcgacccacgg TAGGGCGAATTGGAGCTC | Endogenous Tagging fragments |
| TGGT1_254580  (HAP30) | pCas9 | TAAATGGGGATGTCAAGTT acaagaagctggctcgcaga GTTTTAGAGCTAGAAATAGC | Specific sgRNA |
|  | Generic tagging plasmids | L: ccactgctcacggcggtgagaatgccctgaagaagcagtgc GCTAGCAAGGGCTCGG  T: cccccacatttaaagaccgggcacacaagaagctggctcgc TAGGGCGAATTGGAGCTC | Endogenous Tagging fragments |
| TGGT1_278660  (HAP31) | pCas9 | TAAATGGGGATGTCAAGTT atgcgtgcagaaccgactga GTTTTAGAGCTAGAAATAGC | Specific sgRNA |
|  | Generic tagging plasmids | L: agagagcagctctgacttccaaagcgccggcgattatggcc GCTAGCAAGGGCTCGG  T: gataagagtcgaaagaaaccccgaatgcgtgcagaaccgac TAGGGCGAATTGGAGCTC | Endogenous Tagging fragments |
| TGGT1_313020  (HAP32) | pCas9 | TAAATGGGGATGTCAAGTT ggaaggggtgctcacgtcag GTTTTAGAGCTAGAAATAGC | Specific sgRNA |
|  | Generic tagging plasmids | L: gtgacgtcgctgagcaggtcgcgcatgcagacagagacgcg GCTAGCAAGGGCTCGG  T: agaacacagaatggaacccttttaggggaaccacccccctg TAGGGCGAATTGGAGCTC | Endogenous Tagging fragments |
| TGGT1_209940  (HAP33) | pCas9 | TAAATGGGGATGTCAAGTT gctttaacatcagacgagcg GTTTTAGAGCTAGAAATAGC | Specific sgRNA |
|  | Generic tagging plasmids | L: aattgccagcgaaaaactctggctgtaccaagtcttctcta GCTAGCAAGGGCTCGG  T: ggcgccagcaaaatcaacactcctcagccagcgagccgcgc TAGGGCGAATTGGAGCTC | Endogenous Tagging fragments |
| TGGT1_208050  (HAP34) | pCas9 | TAAATGGGGATGTCAAGTT ggtggacaagagacctgggg GTTTTAGAGCTAGAAATAGC | Specific sgRNA |
|  | Generic tagging plasmids | L: aactttatcatgccggcaccacggagggccaagagcgagag GCTAGCAAGGGCTCGG  T: gcggatgattgatgacccatttggggtggacaagagacctg TAGGGCGAATTGGAGCTC | Endogenous Tagging fragments |
| TGGT1_230420  (HAP35) | pCas9 | TAAATGGGGATGTCAAGTT gaagagactgccgagacgcg GTTTTAGAGCTAGAAATAGC | Specific sgRNA |
|  | Generic tagging plasmids | L: tgagcctgtcgcgctcccagtcgcagctgcgcaagctgcag GCTAGCAAGGGCTCGG  T: gggaaaaaggaaaactcgacatttcgaggcgccttccacgc TAGGGCGAATTGGAGCTC | Endogenous Tagging fragments |
| TGGT1_304750  (HAP36) | pCas9 | TAAATGGGGATGTCAAGTT acaggagccgtcaacggctg GTTTTAGAGCTAGAAATAGC | Specific sgRNA |
|  | Generic tagging plasmids | L: gagcagtcgtcactgcaacagtgaaagatgtgaaaaagtct GCTAGCAAGGGCTCGG  T: ttgcgaatctcgcaacggcagcatgcgaaaatgaaccccag TAGGGCGAATTGGAGCTC | Endogenous Tagging fragments |
| TGGT1_270220  (HAP37) | pCas9 | TAAATGGGGATGTCAAGTT gagtctggtagtacgacctg GTTTTAGAGCTAGAAATAGC | Specific sgRNA |
|  | Generic tagging plasmids | L: tcttaggtcccctgttttttggctgttacgttacatggact GCTAGCAAGGGCTCGG  T: tagtttgattttaaacagatcaaaaacggcaacctcctcag TAGGGCGAATTGGAGCTC | Endogenous Tagging fragments |
| TGGT1_262610  (HAP38) | pCas9 | TAAATGGGGATGTCAAGTT gttcgaactaggtacgggta GTTTTAGAGCTAGAAATAGC | Specific sgRNA |
|  | Generic tagging plasmids | L: tgggagctggagcctctccgtttatgcgagctagctatcag GCTAGCAAGGGCTCGG  T: acctttcttcttttctacagcgcggagaccttgtcccttac TAGGGCGAATTGGAGCTC | Endogenous Tagging fragments |
| TGGT1_319740  (HAP39) | pCas9 | TAAATGGGGATGTCAAGTT atggtggtgagacttcgccg GTTTTAGAGCTAGAAATAGC | Specific sgRNA |
|  | Generic tagging plasmids | L: cacaaacatacacctttgaaaaacgagttccagtctgttca GCTAGCAAGGGCTCGG  T: cctctgagaaaggcgagttgggtcggtctttccccccacgg TAGGGCGAATTGGAGCTC | Endogenous Tagging fragments |
| TGGT1_316260  (HAP40) | pCas9 | TAAATGGGGATGTCAAGTT atgcgaggctgctcagtgtc GTTTTAGAGCTAGAAATAGC | Specific sgRNA |
|  | Generic tagging plasmids | L: tccgaaagatgcaagagacgttctgtctaaagagttttgac GCTAGCAAGGGCTCGG  T: attgctttctcgacggacccgaagaatggaacatgccagac TAGGGCGAATTGGAGCTC | Endogenous Tagging fragments |
| TGGT1_266750  (HAP41) | pCas9 | TAAATGGGGATGTCAAGTT agccgacatggagtcctcaa GTTTTAGAGCTAGAAATAGC | Specific sgRNA |
|  | Generic tagging plasmids | L: tgcaggaagatggggaggaaaagaagaaggtggagcggcaa GCTAGCAAGGGCTCGG  T: tacaccacacgcgacagaaaatgcagccgacatggagtcct TAGGGCGAATTGGAGCTC | Endogenous Tagging fragments |
| TGGT1_233760  (HAP42) | pCas9 | TAAATGGGGATGTCAAGTT gaacaccatcagcaggtctg GTTTTAGAGCTAGAAATAGC | Specific sgRNA |
|  | Generic tagging plasmids | L: tccccgcctctctgcaggaagcgcatccgtacactggcatt GCTAGCAAGGGCTCGG  T:aagcgagacaccgaggaaagggaaagaacaccatcagcaggTAGGGCGAATTGGAGCTC | Endogenous Tagging fragments |
| TGGT1_307860  (HAP43) | pCas9 | TAAATGGGGATGTCAAGTT gaggcgggagtggtgcggag GTTTTAGAGCTAGAAATAGC | Specific sgRNA |
|  | Generic tagging plasmids | L: gtatgcactgcttagaagaagtttctcatatctggttccag GCTAGCAAGGGCTCGG  T: gttttctatcaaccctatccagtagagtttctcttcccctc TAGGGCGAATTGGAGCTC | Endogenous Tagging fragments |
| TGGT1_281640  (HAP44) | pCas9 | TAAATGGGGATGTCAAGTT aacgcttctggcaaccttct GTTTTAGAGCTAGAAATAGC | Specific sgRNA |
|  | Generic tagging plasmids | L: cgcctctcacgcagacggagctgcggaaactgctttcgcag GCTAGCAAGGGCTCGG  T: ttgcatgtgtatatgcatacatttgaacgcttctggcaacc TAGGGCGAATTGGAGCTC | Endogenous Tagging fragments |
| TGGT1_210380  (HAP45) | pCas9 | TAAATGGGGATGTCAAGTT agagggtgagcggagtgtat GTTTTAGAGCTAGAAATAGC | Specific sgRNA |
|  | Generic tagging plasmids | L: cagcggaaactccctccagcgttcctcactctcgtaccgag GCTAGCAAGGGCTCGG  T: cccgaatccctctctttttcaccgccctcctctctccgata TAGGGCGAATTGGAGCTC | Endogenous Tagging fragments |
| TGGT1_321590  (HAP46) | pCas9 | TAAATGGGGATGTCAAGTT gcatcgtcactagtcgctca GTTTTAGAGCTAGAAATAGC | Specific sgRNA |
|  | Generic tagging plasmids | L: tttcgtggcaagatgtgttcgaagcagaaaactgttccacg GCTAGCAAGGGCTCGG  T: tggcgaacacggacctaaaacacggcatcgtcactagtcgc TAGGGCGAATTGGAGCTC | Endogenous Tagging fragments |
| TGGT1_319550  (HAP47) | pCas9 | TAAATGGGGATGTCAAGTT ggaggacacaaccccaagaa GTTTTAGAGCTAGAAATAGC | Specific sgRNA |
|  | Generic tagging plasmids | L: tgttcgtcctcttcgcgatctttggcgccgtcctagatctg GCTAGCAAGGGCTCGG  T: gcgcggcgcccggaactgcgagagggggaggacacaaccccTAGGGCGAATTGGAGCTC | Endogenous Tagging fragments |
| TGGT1_278960  (HAP48) | pCas9 | TAAATGGGGATGTCAAGTT gtccgatccagctacgaacc GTTTTAGAGCTAGAAATAGC | Specific sgRNA |
|  | Generic tagging plasmids | L: ctgtgtggccggaacaaagcacagagaagcggaaggcagag GCTAGCAAGGGCTCGG  T: gtaaatgcttcttgttggtcttggtgtccgatccagctacg TAGGGCGAATTGGAGCTC | Endogenous Tagging fragments |
| TGGT1_208560  (HAP49) | pCas9 | TAAATGGGGATGTCAAGTT agacttcctacagctgagcg GTTTTAGAGCTAGAAATAGC | Specific sgRNA |
|  | Generic tagging plasmids | L: catctcgttctctaactcgcttatttccagagacagataat GCTAGCAAGGGCTCGG  T: tatttggtgtcttcgcgcaaaaatgagacttcctacagctg TAGGGCGAATTGGAGCTC | Endogenous Tagging fragments |
| TGGT1_278990  (HAP50) | pCas9 | TAAATGGGGATGTCAAGTT acagcgaccatcgacaagag GTTTTAGAGCTAGAAATAGC | Specific sgRNA |
|  | Generic tagging plasmids | L: tgggcatgggaacgaccggcggtggctcagcaaagaaataa GCTAGCAAGGGCTCGG  T: ttgatgcggcgtgaaacaacagcacacagcgaccatcgaca TAGGGCGAATTGGAGCTC | Endogenous Tagging fragments |
| TGGT1_221320  (ACC1) | pCas9 | TAAATGGGGATGTCAAGTT gcagagacacacctctccct GTTTTAGAGCTAGAAATAGC | Specific sgRNA |
|  | Generic tagging plasmids | L: aacgcagggaaacgctcgaaagcgctgctacgccagcctgt GCTAGCAAGGGCTCGG  T: agtcaatgattccggttcatctctgcgcagagacacacctTAGGGCGAATTGGAGCTC | Endogenous Tagging fragments |
| pL-AID-GFP-HXGPRT | pL-AID-3HA-HXGPRT | F: TAACCCGGGCATATGTAGAAA  R: gccagagccggcgcgccca | Flagment 1 |
|  | GFP containing plasmid | F: gggcgcgccggctctggcatgGTGAGCAAGGGCG  R: CTACATATGCCCGGGTTACTTGTACAGCTCGTCCAT | Flagment 2 |
